# Macroecology suggests cancer-causing papillomaviruses form non-neutral communities

**DOI:** 10.1101/187047

**Authors:** Marta Félez-Sánchez, Carmen Lía Murall, Ignacio G. Bravo

**Affiliations:** Infections and Cancer Laboratory, Catalan Institute of Oncology (ICO), Barcelona, Spain; Bellvitge Institute of Biomedical Research (IDIBELL), Barcelona, Spain; Laboratory MIVEGEC, French National Center for Scientific Research (CNRS), Montpellier, France.

**Keywords:** HPV, infection and cancer, species abundance distributions, competition, neutrality, vaccine, type replacement, host-parasite interactions, multiple infections

## Abstract

Chronic infection by oncogenic Human papillomaviruses (HPVs) leads to cancers. Public health interventions, such as cancer screening and mass vaccination, radically change the ecological conditions encountered by circulating viruses. It is currently unclear how HPVs communities may respond to these environmental changes, because little is known about their ecology. Predicting the impact on viral diversity by the introduction of HPV vaccines requires answering the unresolved question of how HPVs interact. Although it is commonly believed that they do not interact (neutral theory), there are suggestions that HPV types may compete for resources or via the immune response (niche-based or non-neutral theory). Here, we applied for the first time established biodiversity measures and methods to epidemiological data in order to assess whether niche-partitioning or neutral processes are shaping HPV diversity patterns at the population level. We find that as infections progress toward cancer, HPVs communities become more uneven and a few HPVs play a stronger dominance role. By fitting species abundance distributions, we found that neutral models were always out-performed by non-neutral distributions, both in asymptomatic infections and in cancers. Our results suggest that temporally moving from a more even to a less even community implies an increase in competition, probably due to environmental changes linked to infection progression. More ecological thinking will be required to understand present-day interactions and to anticipate the future of the long lasting interactions between HPVs and humans.

**SIGNIFICANCE STATEMENT:** Human papillomaviruses (HPVs) are very diverse. Infections by HPVs are very common and chronic infections may lead to cancers. The more oncogenic HPVs are now targetted by effective vaccines, and this has raised the question of whether there may be a viral replacement if these dominant types were removed. This is a medical version of a classical ecological controversy, namely how much biodiversity distributions and community dynamics are explained by neutral theory plays out across ecosystems. For HPVs, epidemiologic studies before and after the vaccination have led to the widespread belief that these viruses do not interact. Here, we apply different methods developed in macroecology to the best available epidemiologic data to address this issue. Consistently, we find that HPVs form non-neutral communities. Instead, competitive niche-partitioning process and dominance explain best HPVs communities. We also find that the vaccine might not change such competitive niche processes. Beyond clinical implications, this garners support that niche processes often best explain biodiversity patterns, even in human viral communities.

## INTRODUCTION

Distribution and abundance of species is a historical and central interest of ecology. One research school has focused on niche-based explanations to address the structuring of communities and the distributions of species across ecosystems (1). Meanwhile, neutral theories have challenged this niche-centrist view, suggesting that random dispersal, ecological drift (*i.e.*, stochasticity in births and deaths) and speciation may better explain ecological patterns (2). Specifically, Hubbell’s *Unified Neutral Theory of Biodiversity and Biogeography* (UNTB) argues that species are ecologically equivalent, *i.e.* functionally similar (3). Both niche processes and neutrality shape metacommunity patterns (*e.g.*, species-area or species-occupancy relationships), and thus, inferring from these patterns what potential underlying processes might have molded them has a long tradition in ecology. Fitting distributions to species abundance curves is a common way to infer such underlying processes, especially since several alternative distributions have been derived from either niche-based or neutral models (4-6). Our knowledge about host-associated microbial communities has grown mature enough to incorporate ecological approaches to elucidate multispecies interactions and dynamics. Here we apply these macroecology methods to try and understand communities of important viruses in the human virome.

Whether viral communities are structured by neutral or niche processes has not been addressed at great length. For certain viruses, lineages strongly compete with one another, so that competitive exclusion occurs rapidly, some at the host-population level (*e.g.* Influenza viruses (7)), but also within-host during the course of the infection (*e.g*., *Hepatitis C virus* (8) or *Human immunodeficiency virus* (9, 10)). Indeed, at the cellular level, many different viruses have evolved mechanisms to hinder the entry of new virions into already infected host cells thus preventing from sharing intracellular resources, and precluding recombination between viral strains (*e.g.*, in *Alphaherpesviruses* (11)). In this study, we will focus on Papillomaviruses (PVs), a large family of viruses, with hundreds of stable and largely divergent viral linages (known as “types” in the field; often considered as the relevant taxonomic unit of study at the genotypic and phenotypic level) that are known to coexist with each other inside the same individual hosts, often for years.

PVs are a numerous family of small dsDNA viruses that infect virtually all mammals, as well as other aminotes and fishes (12). Human PVs (HPVs) are the most studied members in the family, because of their medical importance. A handful of closely related HPVs are responsible for around one third of all human cancers linked to infections (13), and HPVs are the causative agents of cancers of the cervix, vagina, anus, vulva, penis and head and neck (14). However, oncogenic HPVs are a clear minority among the more than 200 different HPVs hitherto described: most HPVs are retrieved from healthy skin and mucosas and are never found associated to lesions; some HPVs cause benign wart-like lesions; and only a few can be classified as oncogenic (see (12) and the Papillomavirus Episteme https://pave.niaid.nih.gov).

Public health interventions to decrease the burden of HPVs-associated cancers include systematic screening to detect HPVs chronic anogenital infections *-e.g.* the Papanicolau test- and more recently mass vaccination against the most oncogenic HPVs. The introduction of human-driven selective pressures targeting a subset of the circulating PV diversity implies an important change in the ecological pressures to virus circulation (12). Therefore, evolutionary and ecological considerations of vaccines and PV dynamics have both fundamental and clinical implications. It is currently unclear how viral communities might respond to these environmental changes. This is often referred to in the PV field as the *type replacement problem,* the underlying question being whether upon a hypothetical disparition of the highly oncogenic HPVs targeted by vaccination their niche (in terms of oncogenicity and cancer burden) will be occupied by other HPVs. Predicting the impact caused by the introduction of HPVs vaccines on viral diversity requires an answer to the still unresolved question of interactions between HPVs.

Little is known about interactions between HPVs. The most common hypothesis for interactions between HPVs is simply that they do not interact (15-18). Studies using classic statistical methods have concluded that infections by HPVs occur randomly and lead to cervical disease independently, so that type replacement after vaccination is unlikely (16-20). However, theoretical works at both the within-host (21) and epidemiological levels (22-24) have supported the idea that HPVs infecting the same host likely interact with one another *via* the immune system (both innate and adaptive). Most vaccine trials have not detected significant increases in prevalence of non-vaccine HPVs in vaccinated healthy women (20, 25) which seems to support the neutrality hypothesis. Nevertheless, two HPVs have been flagged as potentially having a competitive advantage (17) and certain non-vaccine types, including probably oncogenic HPVs, displayed higher prevalence in vaccinated patients (26).

Here, we have applied ecological methods to epidemiological data to study which intra-host processes (niche or neutral) are more likely to explain epidemiological *(i.e.,* metacommunity) patterns of HPVs communities.

## RESULTS

### Diversity patterns change with disease progression

Communities of HPVs in the uterus cervix are by far the best described across all stages of the natural history of the disease, from initial asymptomatic infections to invasive cancers. In order to describe cervical HPVs community composition, we performed a correspondence analysis (CA) using viral prevalence data in four different clinical stages and in different geographical regions (see Methods for details). HPVs were stratified based on their carcinogenicity according to the International Agency for Research on Cancer (IARC) classification: carcinogenic HPVs (HPV16, 18, 31, 33, 35, 39, 45, 51, 52, 56, 58, 59); probably or possibly carcinogenic HPVs (HPV26, 30, 34, 53, 66, 67, 68, 69, 70, 73, 82, 85, 97); these two groups are often referred to in the literature as “high-risk” HPVs; finally, HPVs not classifiable as to their carcinogenicity to humans, typically referred in the literature as non-oncogenic or “low-risk” HPVs (HPV6, 11, 44, 74, 7, 40, 91, 57, 81, 29). Our results (Figure 1A) are consistent with the current understanding of cervical disease as they clearly show that HPVs classified as non-oncogenic are more strongly associated with viral communities in asymptomatic and low-grade lesions, whereas oncogenic HPVs are mainly associated to high-grade and cancer communities.

**Figure 1:**
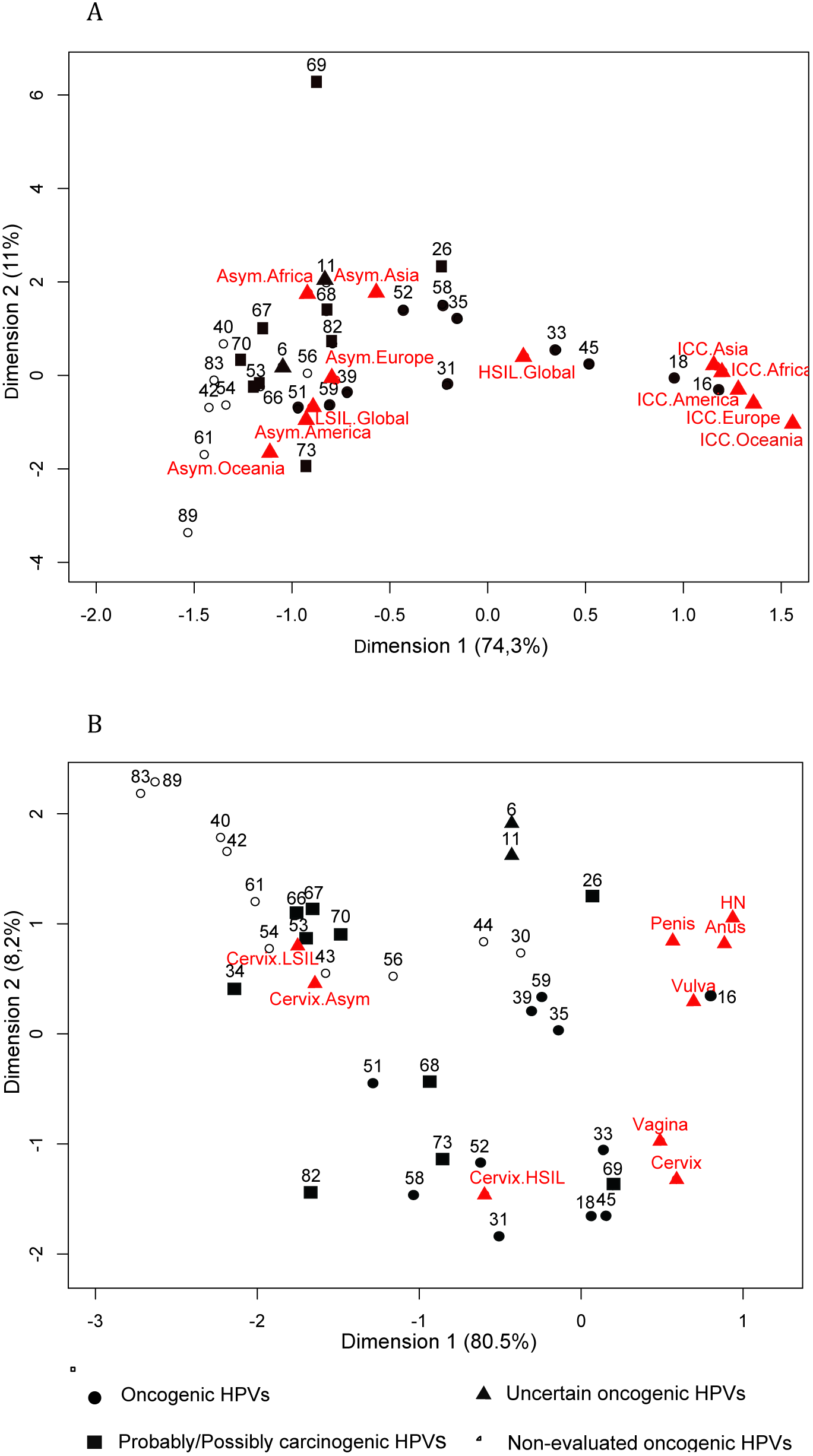
**Correspondence analysis using type prevalences for communities of HPVs.** A) Includes data for cervical communities stratified for each continent in asymptomatic (Asym), low-grade lesions (LSIL), high-grade lesions (HSIL) and invasive cancer (ICC) communities. B) Includes different cancer anatomical locations: cervix, vagina, vulva, anus and head and neck (HN). Full circles, oncogenic HPVs (group 1 IARC classification); squares, probably/possibly oncogenic HPVs (group 2A/2B IARC classification); triangles, uncertain oncogenic HPVs (group 3 IARC classification); empty circles, non-evaluated oncogenic HPVs (non-evaluated by IARC)

While HPVs can infect various anatomical regions of the body and are associated with a number of cancers, only the uterus cervix is systematically screened for early cancer detection. Our knowledge of the natural history of the infection in other anatomical sites is thus poorer and the quality and amount of data about HPVs in asymptomatic infections or in pre-neoplastic lesions also lag behind. Consequently, for the vagina, vulva, anus, penis, and head and neck we only have reliable data for HPV prevalence in cancers. The results of our CA including these cancer communities (Figure 1B) showed that, similar to cervical cancers, oncogenic HPVs were found to be strongly associated to these cancer communities (Figure 1B). Remarkably, we found that HPV26 (classified as probably/possibly carcinogenic) was strongly associated with other cancer communities different to the cervix, mainly due to the high prevalence of HPV26 in head and neck cancers.

### Communities of HPVs become more uneven with advancing disease progression

We have analysed the diversity profile of HPVs communities across the different stages of the cervical infection leading to cancer (Figure 2A), as well as for the different cancers associated to HPVs infections (Figure 2B). A single community was defined as the assemblage of HPVs recorded at each stage of the infection in each anatomical region. Diversity was proxied using the diversity index (*q*) at different orders: species richness (*q*=0), Shannon entropy (*q*=1), and the inverse of Simpson index (*q*=2). The more uneven the distribution of relative abundances, the more steep the diversity profile declines with the increasing order of the diversity index. We observed first that all cervical communities exhibited similar levels of HPV species richness (*q*=0), independently of the stage of progression. For higher orders of the diversity index we observed a decline with the progression of the disease, indicating that in advanced stages of the infection (cancer and high-grade lesions), viral communities become less even in comparison with early stages (asymptomatic and low-grade lesion). HPVs communities in asymptomatic and low-grade lesions were moderately uneven while high lesion grades and cancer displayed the steepest slopes and thus were highly uneven communities (Figure 2A, Table S1). Since the number of different HPVs is not different in the various communities, we interpreted that the observed changes in evenness during cancer progression are likely due to changes in the species interaction dynamics.

**Figure 2:**
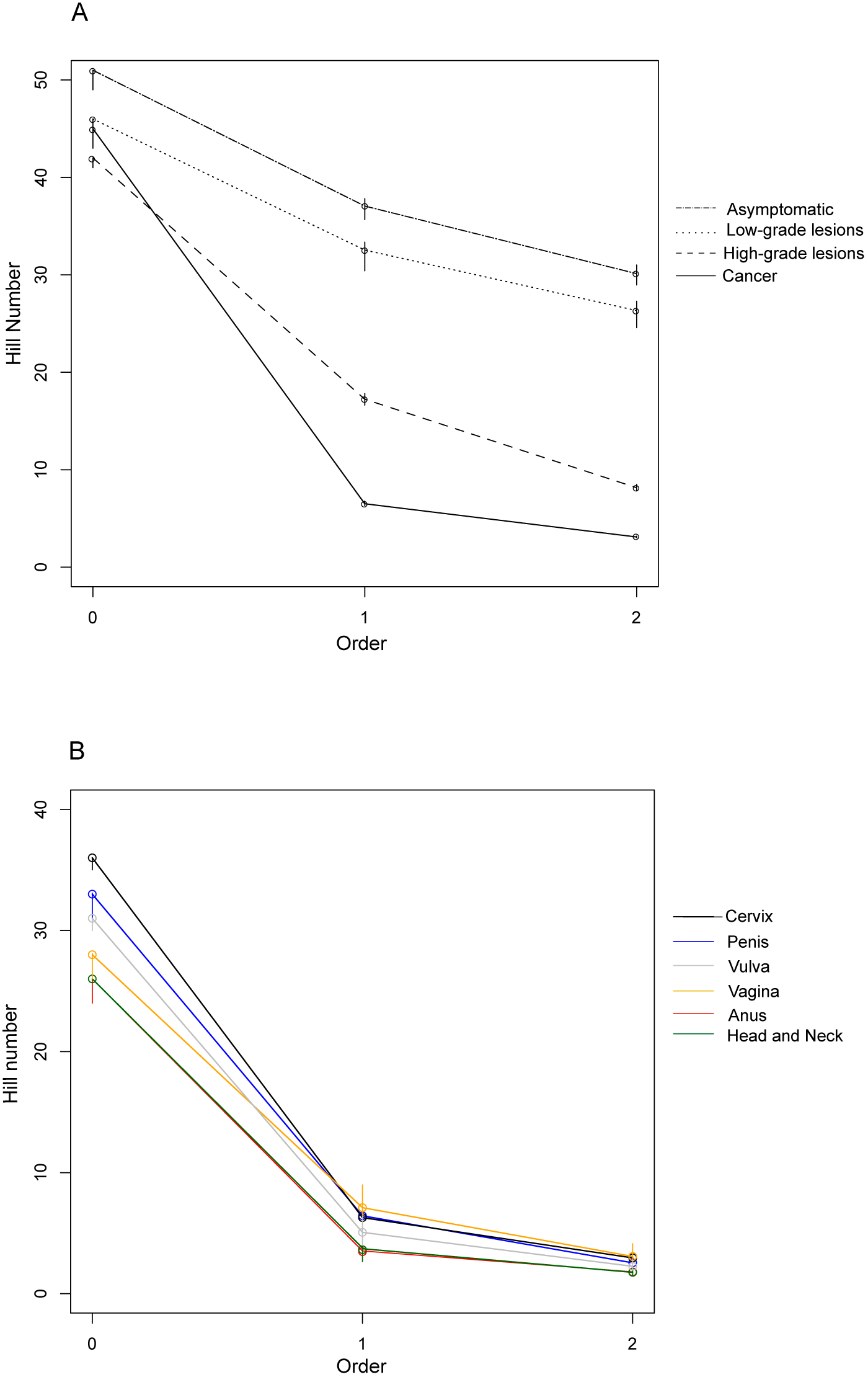
**Diversity (*^q^D*) profiles for A) different stages of HPV infection in cervical lesions (Asymptomatic, Low-grade Lesions, High-grade Lesions and cancer) and B) squamous carcinomas of different anatomical regions (cervix, vulva, vagina, anus, penis and head and neck).** Three orders of diversity (*q* = 0, 1 and 2) were calculated at each stage of the infection. As parameter *q* increases, rare types are weighted less and *q* becomes a measure of evenness.

We also found that the species richness of HPVs in cervical cancer communities was highest compared to other cancers (Figure 2B, Table S1), whereas head and neck and anal cancer communities were the least rich. These differences in terms of diversity were sharply reduced with increasing values of *q*. Overall, communities of HPVs in infection-related cancers of various anatomical regions were highly uneven communities with *q*=2 ranging from 1.8 to 3, demonstrating that only a few HPVs completely dominate cancer communities.

### Very few types dominate HPV communities

In order to improve the ecological understanding of HPVs, we have also studied the species abundance distribution (SAD) of each viral community. We represented these distributions using rank-abundance plots, which were generated after fitting different alternative SADs to the data. Rank-abundance distributions (RADs) were drawn by representing HPV rank in a descending order on the abscissa and the corresponding HPV frequencies on the ordinate axis. We confronted the following five alternative models with each data set to assess how well each accounted for the data: Broken-stick (27), geometric distribution (28), Power law (29), Poisson lognormal (30), and the Zero-sum multinomial (3) (Table S2). We assessed robustness of the results by data resampling (see Methods for more details). This resampling strategy was applied to all anatomical sites and in the case of the cervix separately to all stages in the natural history of the infection and for data from asymptomatic infection in the cervix after stratifying by geographical origin and by age.

For communities of HPVs in asymptomatic infections and in low-grade lesions, the geometric distribution and the broken-stick model displayed the lowest AICc values (with ΔAICc<3 between them) compared with the other candidate distributions (Figure 3, Table S3). Overall, the best fits for communities in the early stages of the natural history of the disease corresponded to a niche-partitioning model (broken-stick or geometric) or to independence of interactions between HPVs (geometric). In the cases in which broken-stick and geometric were tied, the lognormal distribution (compatible with non-neutral interactions between HPVs) systematically outperformed the neutral model ZSM (compatible with neutral interactions between HPVs). Thus, in the cases of such ties for conflicting models, it is more likely that niche-partitioning and not neutral processes underlie these communities. Overall, our results suggest that during early stages of the natural history of the HPV infection, niche-partitioning processes structure communities of HPVs, but create a population level pattern that can be difficult to distinguish from neutrality. We found similar results across geographical regions and ages (see Table S3 in Supplementary Material).

**Figure 3:**
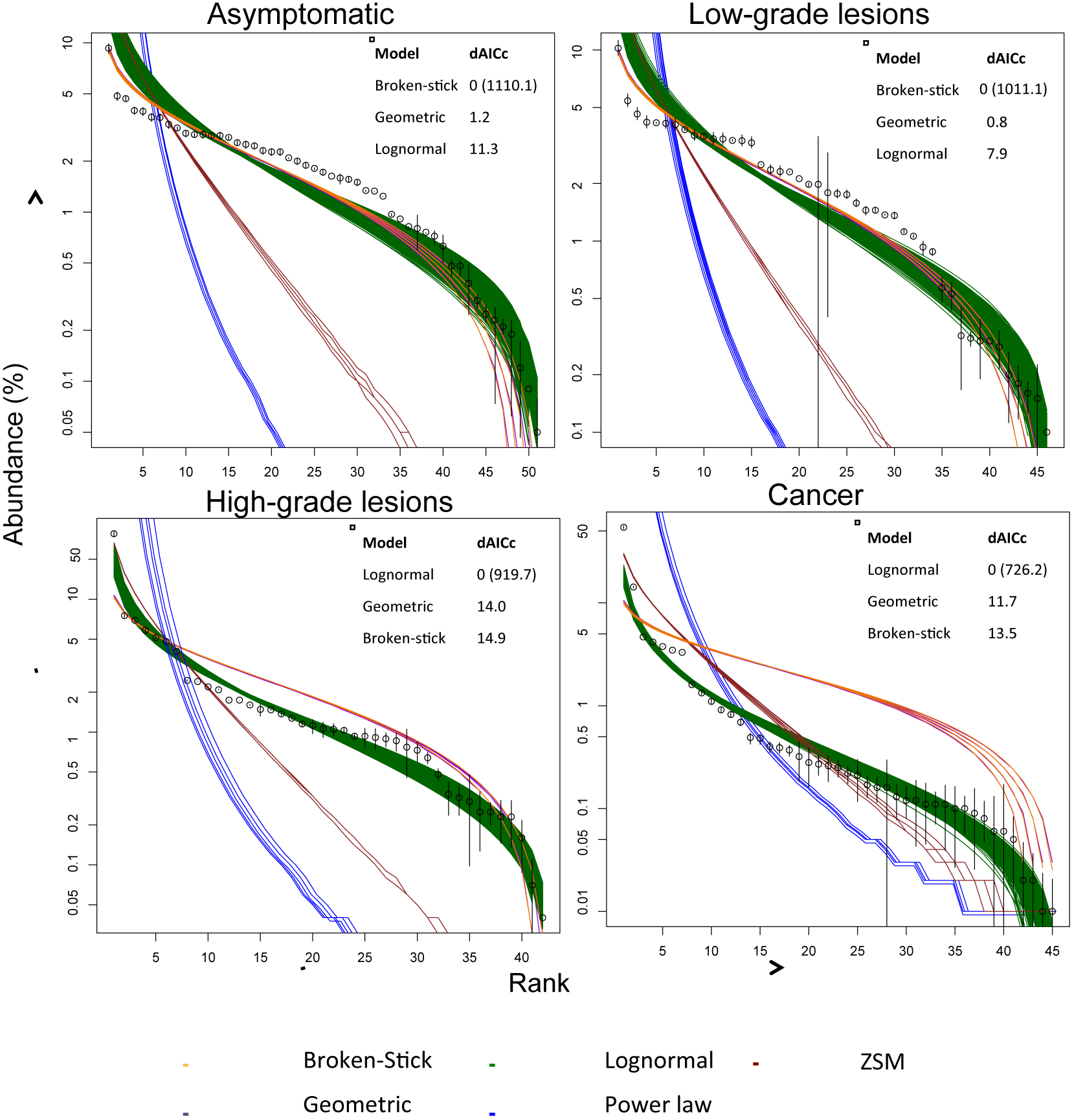
**Rank abundance distribution for cervical communities of HPVs at different stages of cervical infection**. A) Asymptomatic (N=86,696); B) Low-grade Lesions (N=46,402); C) High-grade lesions (N=51,616) and D) Invasive Cancer (N=50,084). Model fits are also shown for the 1000 random communities resampled from empirical data. Fits are shown for 5 models: Broken-Stick (orange), Lognormal (green), Geometric (violet), Power law (blue) and Zero Sum Multinomial (brown). The inset shows the calculated Akaike Information Criterion for the best fit (in brackets), and the difference for the second and third best-fits. Broken-stick curve is not visible as it overlaps with the geometric model one.

For communities in high-grade lesions and in cervical cancer the best fit corresponded always to the lognormal model (Figure 3, Table S3). This was also the case for 100% of the randomly generated communities. The lognormal distribution also fitted best all cancer communities, independently of anatomical location (Figure 4, Table S3). Only in one community was the lognormal statistically tied to another distribution (the ZSM, ΔAICc=2.8) and that was in cervical carcinoma using a more restricted dataset (Figure 4, Table S3). However for this same community in the meta-analysis dataset the ZSM was out-performed by the lognormal and by other distributions (Figure 4). Except for this one tie, the ZSM was systematically outperformed by the lognormal in all our datasets. Thus, in the late stages of the natural history of the infection, viral communities are structured by non-neutral interactions between HPVs.

**Figure 4:**
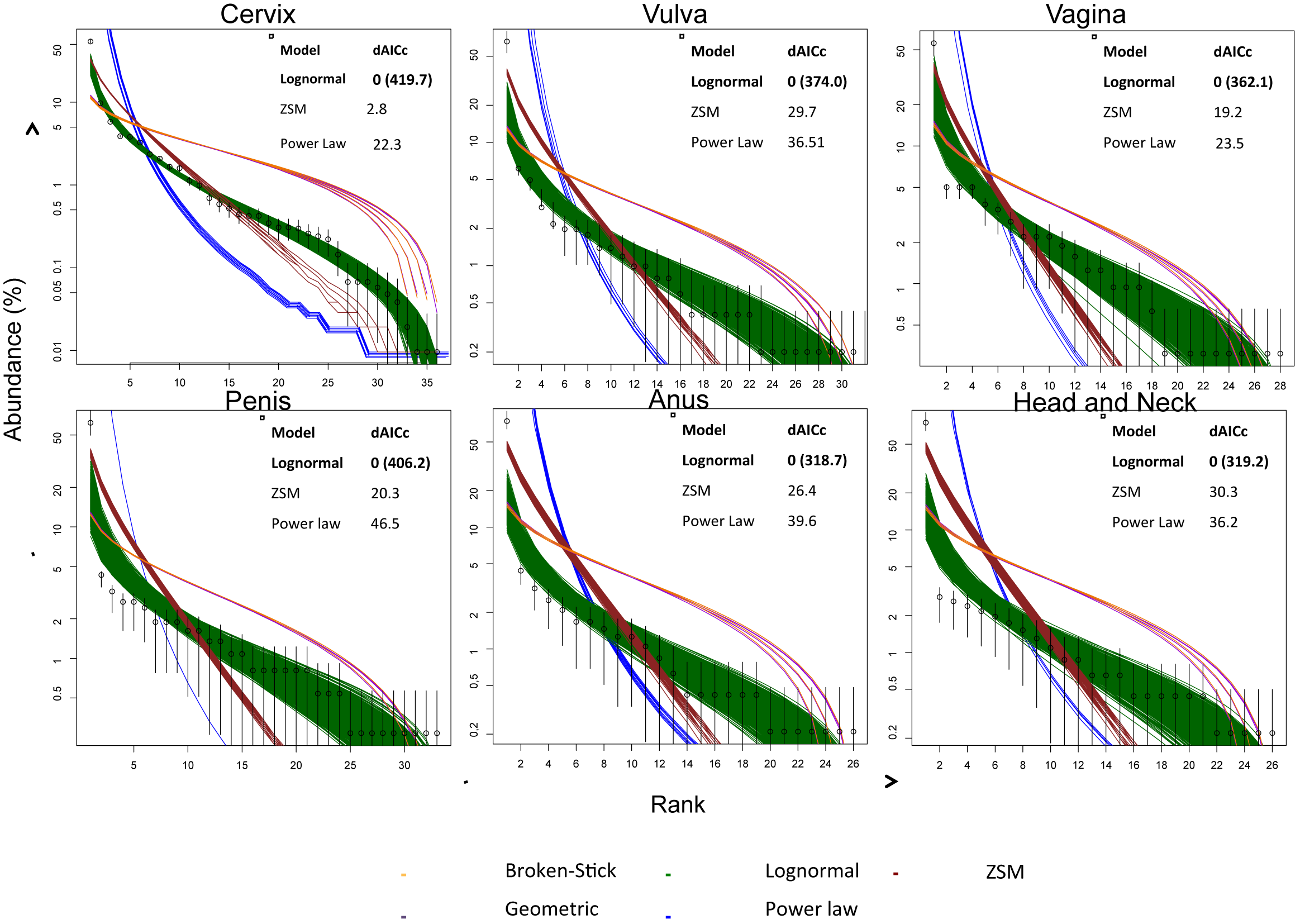
**Rank abundance distribution for cancer communities at different anatomical locations.** A) Cervix (N=8,977); B) Vulva (N=488); C) Vagina (N=303), D) Penis (N=334), E) Anus (N=438), Head and Neck (N=453) Model fits are also shown for 1,000 random communities resampled from empirical data. Fits are shown for 5 models: Broken-Stick (orange), Lognormal (green), Geometric (violet), Power law (blue) and Zero Sum Multinomial (brown). The inset shows the calculated Akaike Information Criterion for the best fit (in brackets), and the difference for the second and third best-fits. Note the very large gap for the first and the second more frequent HPV. Broken-stick curve is not visible as it overlaps with the geometric model one.

Remarkably, the strong drop in prevalence between the first ranked HPV type and the rest (see Figures 3 and 4) was not captured well by any of the theoretical model distributions used. This gap was especially large in the cancer data in all anatomical locations, where HPV16 largely dominates other HPVs, particularly for head and neck cancers. Thus, all models (even those assuming non-neutral interactions) anticipate more evenness in these first few ranked types than it is actually observed. This failure of all models at capturing the steep drop at the beginning of the curve, suggests that strong dominance of HPV16, combined with the long tail of rare HPVs, are not easily captured by current SAD models.

### Interactions among HPVs may remain unchanged post-vaccination

We have also fitted SADs to the few available pre- and post-vaccination asymptomatic cervical data, in order to assess the potential impact of vaccines on communities of HPVs. We found that the best-fitting SAD models for both communities remained the geometric distribution and the broken-stick model (followed by the lognormal distribution with ΔAICc<9 in both cases), suggesting that the novel vaccination pressures might not change the processes driving type distribution in HPV communities (Figure 5, Table S3). Therefore, even if the vaccine reduced the prevalence of the targeted viruses, which often dominate the community, interactions in the viral community in asymptomatic infections may remain sufficiently uneven to reject neutrality.

**Figure 6:**
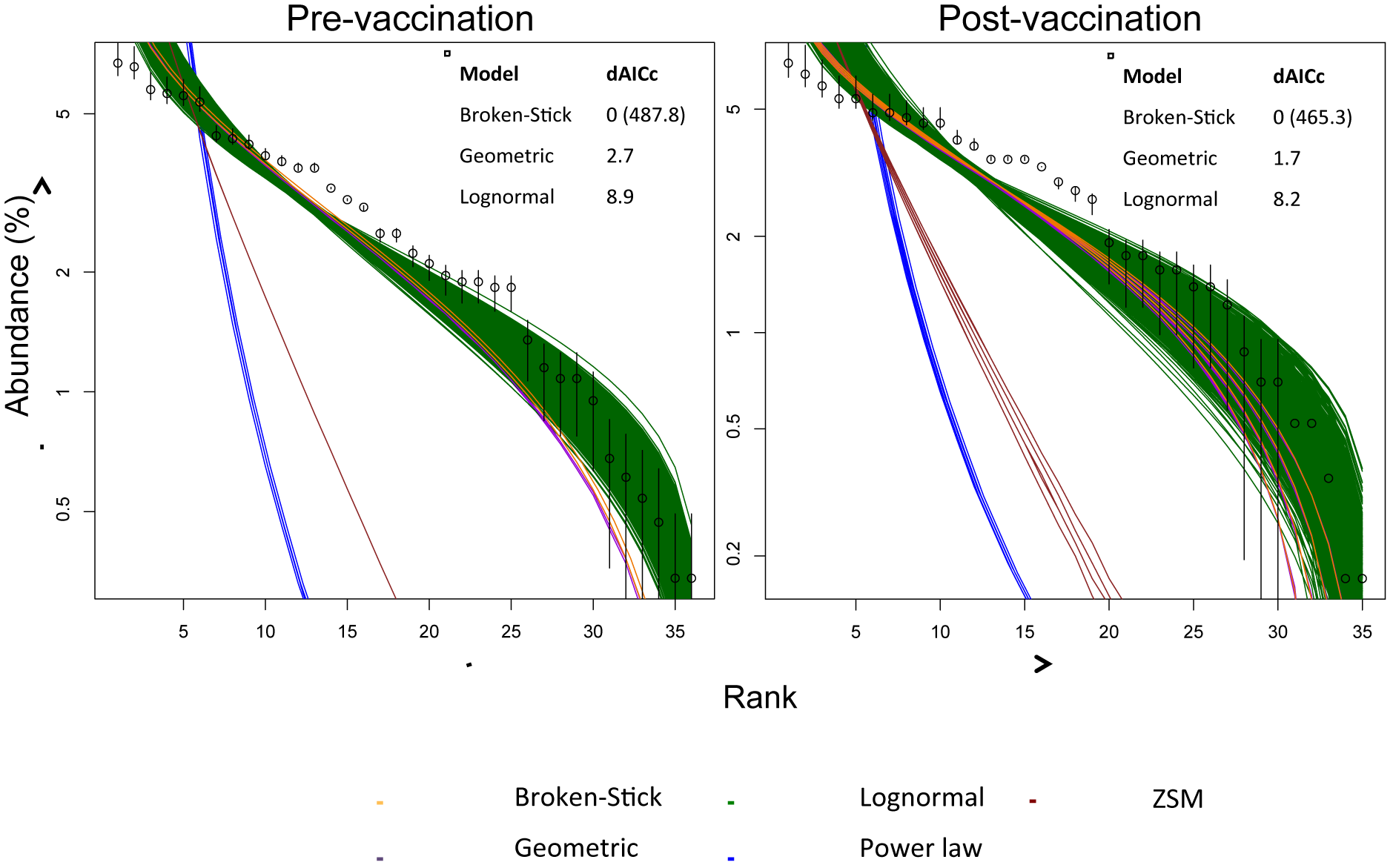
**Rank abundance distribution for A) pre-vaccination (N= 1479) and B) post-vaccination (N=575) asymptomatic cervical HPV communities**. Model fits are also shown for the 1000 random communities resampled from empirical data. Fits are shown for 5 models: Broken-Stick (orange), Lognormal (green), Geometric (violet), Power law (blue) and Zero Sum Multinomial (brown). The inset shows the calculated Akaike Information Criterion for the best fit (in brackets), and the difference for the second and third best-fits. Broken-stick curve is not visible as it overlaps with the geometric model one.

## DISCUSSION

We analyse here the diversity patterns of HPVs along the gradient of clinical presentations of infection, from asymptomatic to cancer. With the ultimate aim of understanding the nature of interaction dynamics in oncogenic pathogen communities, we specifically tried to discern between neutral and non-neutral viral interactions in HPV communities.

First, we show the sharp distinction between communities of HPVs in health and disease. On one hand, we confirm the clear gradient in HPV repertoire and prevalence (Figure 1), with HPVs classified as non-oncogenic being more strongly associated to viral communities in asymptomatic and low-grade lesions, and oncogenic HPVs mainly associated to high-grade and cancer communities. On the other hand, we uncover the transition from an evenly distributed community to a highly uneven community along the natural history of these infections (Figure 2). We find that species richness changes little with disease progression while dominance clearly increases, suggesting that there is an increase in the strength of the interactions of the types.

Second, our analyses of global data on viral prevalence in different anatomical sites using macroecology methods show the low explanatory power of neutrality to account for viral community diversity (Figure 3 and Figure 4). We find that interactions in the initial steps of the carcinogenic process are best fitted by a niche partitioning model, while those in the advanced steps correspond best to dominance, non-neutral interaction model.

Given the natural history of infections by HPVs, niche partitioning appears biologically meaningful in the initial steps of the infection. The target cells for HPVs are located in the basal layer of the epithelium, and virions reach them through epithelial abrasions and microtroauma. Since the infection targets are not constantly exposed and available for viral entry, different viruses compete for access to basal cells through these entries of the epithelium. At any one time the availability of the target cell resource is thus limited and dependent on the stochastic opening up of abrasions.

Previous epidemiological studies of HPVs prevalence in non-cancer communities have used statistical approaches so that if the probability of finding a pair of HPV types together in a co-infection is lower than expected by chance, it is then assumed that these two HPVs compete. The most common of these approaches is odds ratios (reviewed in (16)), while others have used logistic regression models *(e.g.* (31)). Recently a hazard ratios approach was used on cohorts of women, where HPV acquisition or clearance was analysed with Cox proportional hazards regression (18). All of these studies have generally found that pairwise co-occurrence patterns do not significantly deviate from independence. This appearance of independence is also found by our study in the asymptomatic and low-grade lesion HPV communities (but not in high-grade lesions nor cancer) due to the statistical tie in AICs that we find between the broken-stick and the geometric distribution. Our results demonstrate the difficulty in inferring local processes from global patterns, when both independence and niche processes can create similar patterns (32-34). The broken-stick (niche partitioning model) and the geometric distribution (statistical independence model) are derived from completely opposite sets of assumptions and yet they are, in fact, related because the geometric distribution is the discrete equivalent of the exponential distribution which was shown to generate the same RAD as the broken-stick (28). In this study, we argue that, given that the Poisson lognormal systematically outperformed the ZSM in all cases, niche partitioning is more likely than neutrality as a explanatory scenario for viral interactions in the early stages of the HPVs infection (independence being a special case of neutrality). This result matches also similar studies for other ecosystems, where neutrality performs poorly against the lognormal (5, 35, 36).

The movement towards a lognormal distribution with disease progression, coupled with reduced community evenness suggests an increase in competitive ability of some types in the community during carcinogenesis. Disease progression towards cancer is accompanied by a change in the local environment, namely an increase in immunity activity *(e.g.,* more immunity effectors, cytokines, etc.) and the types persisting are using more of the host resources *(i.e.,* leaving less resource and space for other types to invade). Viral loads have been shown to increase with disease progression (37) and it is possible that this may increase competition for newly open abrasions in the mucosa. Relatedly in other anatomical sites, the biology of the site may play a role in increasing the strength of the dominance of the oncogenic types (*e.g.*, head-and-neck and anal cancers have the lowest inverse Simpson index). It is not known whether this could be due to less available resources or to a more active immune response (*e.g.*, being close to lymph nodes) in these particular body sites. It should be noted that this potential increase in competition is not strong, because if this were the case the high-grade and cancer communities would be fitted best by a power law or by other less even SADs, which is not the case.

Post-vaccination data were best fitted by the same distribution as the pre-vaccination data, suggesting then that the vaccine is not changing the underlying processes that govern HPV type distributions in healthy women. While the exact kind or relative strengths of interactions between types have yet to be evaluated, there is a need to unravel the complex interactions between types (direct or indirect) in order to better evaluate the effect of the vaccines long-term. It is important to highlight that the dataset used to perform this analysis was collected over only three years after vaccine implementation, and are from samples across the USA where vaccination coverage is very heterogeneous and often very low in some areas. Thus, these conclusions should be seen as preliminary at best. To evaluate potential type replacement in the post-vaccine era will require other forms of type-specific studies of non-cancer communities that are sufficiently long-term to distinguish true changes in type prevalence from natural fluctuations or noise and that can take into account sampling biases from detection methods (*e.g.*, the unmasking effect; (16)).

In order to check the validity of our results and rule out the possibility of sampling artefacts in our analyses, we checked the effect of the sample size in the selection of the best-fit model (Figure S1). We found that in undersampled communities, the probability to best-fit a Power law model increases. Indeed, the lognormal distribution is believed to better-fit large datasets. Previous published studies showed that in about 500 SADs, the best fit SAD model changed depending on whether the community had been fully censored or not [35]. These study also found that “fully censored” communities were best fit by the lognormal, while “incompletely sampled” communities were best fit by the Power Law model [35]. We also found that the lognormal best described even the smallest datasets. Hence, we can conclude that lognormal fits are not a sampling artefact.

To our knowledge, the only other examples of host-associated microorganism communities that have been analysed for niche and neutral patterns using similar methods are bacterial. Cobey and Lipsitch (38) found that both competition and neutral processes, mainly through immunity interactions, best explained *Streptococcus pneumoniae* diversity patterns across observed carriage populations. Jeraldo and coworkers (39) investigated the roles of niche and neutral processes in shaping gastrointestinal microbiome communities. Using their own method that combines ecological measures with phylogenetic analysis, they found that while community RADs were explained well by the ZSM, further analysis of genomic data was consistent with niche selection as the dominant process. This, once again demonstrates that apparently neutral-like patterns can be generated by underlying non-neutral processes. Bacterial and viral communities have very marked differences so more studies into the roles of niche and neutral processes shaping viral communities are needed.

One possible future direction would be to assess whether non-cancer communities of HPVs could be described by emergent neutrality theory (40-42), where patterns that appear neutral are the outcome of competition and evolutionary processes, such that species evolve to be functionally similar and thus coexist. Most of the HPVs datasets we examined appear as though they could be explained by multimodal SADs, possibly the two-mode Poisson lognormal distribution, which have been found to explain several empirical datasets (42). Indeed, *Alphapapillomaviruses* infecting the mucosa are in many respects functionally similar with only a few clear life history differences in the functions of the oncogenes of oncogenic and non-oncogenic viruses (43). Moreover, as discussed previously (21), data on competition experiments under varying conditions are still needed that to best characterize HPV type interactions. As environmental conditions change, the relative importance of niche and neutral processes can shift and are not static. In addition, it has been well documented that large-scale diversity patterns can be limited in their ability to infer local interactions or to tell how they will change when perturbed (34). Given this and that the PVs are a highly diverse family of viruses that readily coexist, signals of underlying interactions maybe be subtle. Therefore, we call on the HPV community to not be over confident in predictions generated by methods not designed to infer ecological interactions and, instead, seize the opportunity to investigate the interesting ecology of these viruses more mechanistically.

## MATERIALS AND METHODS

### Dataset

For this study the unit of analysis is the HPV at the level of type, according to the definition of the International Committee for the Taxonomy of Viruses, *i.e.*, when we write “different HPVs” we mean “different types of Human Papillomaviruses”. For time-trends analyses, worldwide HPV type-specific prevalence values in different stages of the natural history of the cervical HPV infection (*i.e.* asymptomatic, low-grade squamous intraepithelial lesions (LSIL), high-grade squamous intraepithelial lesions (HSIL) and invasive cervical carcinoma (ICC)) were obtained from a meta-analysis previously published (44) (Table S4). For site-specific analyses, worldwide HPV type-specific prevalence values in squamous carcinomas of different anatomical locations (*i.e.* cervix, anus, penis, vulva, vagina and head and neck) were obtained from the retrospective cross-sectional study by the Catalan Institute of Oncology (45-50) (Table S5). We also obtained information on HPV prevalence values stratified by continent for asymptomatic and invasive carcinomas of the cervix from the WHO/ICO Information Center (https://hpvcentre.net) (Table S6 and S7). Finally, HPV type prevalence values both pre- and post-vaccination asymptomatic cervical communities were obtained from a published study (51) (Table S8). Communities were defined as the species assemblages recorded at each stage of the infection in each anatomical region.

### Correspondence Analysis

HPV type-specific prevalences were subjected to dimensionality reduction techniques to analyse HPV communities on the basis of their type composition. Correspondence analysis (CA) consists in a multivariate statistical method widely used to summarize the lack of independence between objects represented through rows and columns of a matrix (here communities and HPV type prevalences, respectively) as a small number of derived variables, called axes. By definition, the axes are ordered according to the amount of variance in the data explained by them. Data were plotted on the first two axes with the information on the amount of variance explained in these two dimensional representations.

### Non-parametric measures of diversity

We used Hill numbers to estimate diversity within each community. Hill numbers were computed using the following equation for *q* ≠ 1(52).

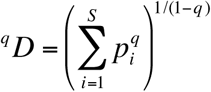

where *S* is the number of species sampled, *p*_*i*_, is the species frequency of the *i*_th_ species, and the parameter *q,* which is the order of diversity, determines its sensitivity to species frequencies. To calculate diversity within each community, *p* was calculated from the vector of the relative abundance of *N* species. As *q* increases, the ^*q*^*D* is more strongly affected by the abundances of the most dominant species and rare species are weighted less. The Hill numbers are directly related to commonly used indexes. When *q =* 0, *^0^D* is equivalent to species richness and when *q =* 2, ^*2*^*D* equals the inverse Simpson index. The measure is undefined when *q =* 1, but the limit as *q* approaches 1 equals the exponential of Shannon entropy.

We chose to use this diversity measure because it quantifies the effective number of species in the dataset, which refers to the number of equally abundant species necessary to produce the observed value of diversity. Moreover, these effective numbers obey the replication principle, which states that if two communities with X equally abundant species are combined, then the diversity of the combined community should be twice that of the original communities (53). Moreover, the unit of these diversity indices is the “effective number of species” (ENS) and thus values of diversity are comparable across different metrics. Hill numbers were computed using the *Vegan* package implemented in R (http://www.r-project.org).

### Species Abundance Distribution

We studied the HPV type-specific distribution of each community by using species abundance distribution (SAD) plots. Model fitting was performed using the *sads* package implemented in R and visualized by means of RAD plots. We evaluated six models: Geometric, Zero-sum multinomial, Poisson Lognormal, Weibull and Power law. These models were chosen because they are most commonly considered to be of ecological relevance. The estimated parameters were inferred by maximum likelihood methods. Model diagnostics were computed by using quantile-quantile and percentile-percentile graphs of the observed vs. predicted abundances. Finally, the observed SADs were compared with the hypothetical models using a Bayesian approach: an Akaike Goodness of fit calculation. The lower the calculated Akaike information criterion (AICc) value, the better the fit.

### Random communities

We generated 1,000 random communities, drawing for each HPV a prevalence value within the 95% Confidence Intervals (CI) of the prevalence in the real data. These synthetic communities were used to calculate the 95% CI of Hill numbers. We also used these randomly generated communities to calculate the number of times that each RAD model best fit each community. When differences in the AICc values among models were below 3, these models were considered as equally probable.

## AKNOWLEDGMENTS

We would like to thank Steve C. Walker and Paulo I. Prado for helpful technical discussions.

## BIBLIOGRAPHY

1. Fisher CK & Mehta P (2014) Proc Natl Acad Sci U S A 111, 13111–13116.

2. Matthews TJ & Whittaker RJ (2014) Ecol Evol 4, 2263–2277.

3. Hubbell S (2001) The Unified Neutral Theory of Biodiversity and Biogeography (Princeton University Press).

4. Magurran AE (2004) Measuring biological diversity (Oxford).

5. McGill BJ, Etienne RS, Gray JS, Alonso D, Anderson MJ, Benecha HK, Dornelas M, Enquist BJ, Green JL, He F, et al. (2007) Ecol Lett 10, 995–1015.

6. Matthews TJ & Whittaker RJ (2014) Frontiers of Biogeography 6.

7. Ferguson NM, Galvani AP, & Bush RM (2003) Nature 422, 428–433.

8. Ramirez S, Perez-del-Pulgar S, Carrion JA, Coto-Llerena M, Mensa L, Dragun J, Garcia-Valdecasas JC, Navasa M, & Forns X (2010) J Gen Virol 91, 1183–1188.

9. Anderson RM & May RM (1996) AIDS 10, 1663–1673.

10. Schmidt WP, Van Der Loeff MS, Aaby P, Whittle H, Bakker R, Buckner M, Dias F, & White RG (2008) Epidemiol Infect 136, 551–561.

11. Meurens F, Keil GM, Muylkens B, Gogev S, Schynts F, Negro S, Wiggers L, & Thiry E (2004) J Virol 78, 9828–9836.

12. Bravo IG & Felez-Sanchez M (2015) Evol Med Public Health 2015, 32–51.

13. Forman D, de Martel C, Lacey CJ, Soerjomataram I, Lortet-Tieulent J, Bruni L, Vignat J, Ferlay J, Bray F, Plummer M, et al. (2012) Vaccine 30 Suppl 5, F12–23.

14. Walboomers JM, Jacobs MV, Manos MM, Bosch FX, Kummer JA, Shah KV, Snijders PJ, Peto J, Meijer CJ, & Munoz N (1999) J Pathol 189, 12–19.

15. Dillner J, Arbyn M, Unger E, & Dillner L (2010) Clin Exp Immunol 163, 17–25.

16. Tota JE, Ramanakumar AV, Jiang M, Dillner J, Walter SD, Kaufman JS, Coutlee F, Villa LL, & Franco EL (2013) Am J Epidemiol 178, 625–634.

17. Tota JE, Chevarie-Davis M, Richardson LA, Devries M, & Franco EL (2011) Prev Med 53 Suppl 1, S12–21.

18. Tota JE, Jiang M, Ramanakumar AV, Walter SD, Kaufman JS, Coutlee F, Richardson H, Burchell AN, Koushik A, Mayrand MH, et al. (2016) PLoS One 11, e0166329.

19. Chaturvedi AK, Katki HA, Hildesheim A, Rodriguez AC, Quint W, Schiffman M, Van Doorn LJ, Porras C, Wacholder S, Gonzalez P, et al. (2011) J Infect Dis 203, 910–920.

20. Palmroth J, Merikukka M, Paavonen J, Apter D, Eriksson T, Natunen K, Dubin G, & Lehtinen M (2012) Int J Cancer 131, 2832–2838.

21. Murall CL, McCann KS, & Bauch CT (2014) J Theor Biol 350, 98–109.

22. Durham DP, Poolman EM, Ibuka Y, Townsend JP, & Galvani AP (2012) J Infect Dis 206, 1291–1298.

23. Pons-Salort M, Letort V, Favre M, Heard I, Dervaux B, Opatowski L, & Guillemot D (2013) Vaccine 31, 1238–1245.

24. Peralta R, Vargas-De-Leon C, Cabrera A, & Miramontes P (2014) Comput Math Methods Med 2014, 542923.

25. Wheeler CM, Castellsague X, Garland SM, Szarewski A, Paavonen J, Naud P, Salmeron J, Chow SN, Apter D, Kitchener H, et al. (2012) Lancet Oncol 13, 100–110.

26. Kahn JA, Brown DR, Ding L, Widdice LE, Shew ML, Glynn S, & Bernstein DI (2012) Pediatrics 130, e249–256.

27. MacArthur RH (1957) Proc. Natl. Acad. Sci 43.

28. Cohen JE (1968) The American Naturalist 102.

29. Newman MEJ (2005) Contemporary Physics 46.

30. Bulmer MG (1974) Biometrics 30, 101–110.

31. Vaccarella S, Soderlund-Strand A, Franceschi S, Plummer M, & Dillner J (2013) PLoS One 8, e71617.

32. Walker SC (2007) Theor Popul Biol 71, 318–331.

33. Ruokolainen L, Ranta E, Kaitala V, & Fowler MS (2009) Ecol Lett 12, 909–919.

34. Al Hammal O, Alonso D, Etienne RS, & Cornell SJ (2015) PLoS Comput Biol 11, e1004134.

35. Walker SC & Cyr H (2006) Oikos 116.

36. Connolly SR, MacNeil MA, Caley MJ, Knowlton N, Cripps E, Hisano M, Thibaut LM, Bhattacharya BD, Benedetti-Cecchi L, Brainard RE, et al. (2014) Proc Natl Acad Sci U S A 111, 8524–8529.

37. Depuydt CE, Jonckheere J, Berth M, Salembier GM, Vereecken AJ, & Bogers JJ (2015) Cancer Med 4, 1294–1302.

38. Cobey S & Lipsitch M (2012) Science 335, 1376–1380.

39. Jeraldo P, Sipos M, Chia N, Brulc JM, Dhillon AS, Konkel ME, Larson CL, Nelson KE, Qu A, Schook LB, et al. (2012) Proc Natl Acad Sci U S A 109, 9692–9698.

40. Holt RD (2006) Trends Ecol Evol 21, 531–533.

41. Scheffer M & van Nes EH (2006) Proc Natl Acad Sci U S A 103, 6230–6235.

42. Vergnon R, van Nes EH, & Scheffer M (2012) Nat Commun 3, 663.

43. Doorbar J, Quint W, Banks L, Bravo IG, Stoler M, Broker TR, & Stanley MA (2012) Vaccine 30 Suppl 5, F55–70.

44. Bzhalava D, Guan P, Franceschi S, Dillner J, & Clifford G (2013) Virology 445, 224–231.

45. Alemany L, Cubilla A, Halec G, Kasamatsu E, Quiros B, Masferrer E, Tous S, Lloveras B, Hernandez-Suarez G, Lonsdale R, et al. (2016) Eur Urol 69, 953–961.

46. Alemany L, Saunier M, Alvarado-Cabrero I, Quiros B, Salmeron J, Shin HR, Pirog EC, Guimera N, Hernandez-Suarez G, Felix A, et al. (2014) Int J Cancer.

47. Alemany L, Saunier M, Tinoco L, Quiros B, Alvarado-Cabrero I, Alejo M, Joura EA, Maldonado P, Klaustermeier J, Salmeron J, et al. (2014) Eur J Cancer 50, 2846–2854.

48. Castellsague X, Alemany L, Quer M, Halec G, Quiros B, Tous S, Clavero O, Alos L, Biegner T, Szafarowski T, et al. (2016) J Natl Cancer Inst 108, djv403.

49. de Sanjose S, Alemany L, Ordi J, Tous S, Alejo M, Bigby SM, Joura EA, Maldonado P, Laco J, Bravo IG, et al. (2013) Eur J Cancer 49, 3450–3461.

50. de Sanjose S, Quint WG, Alemany L, Geraets DT, Klaustermeier JE, Lloveras B, Tous S, Felix A, Bravo LE, Shin HR, et al. (2010) Lancet Oncol 11, 1048–1056.

51. Markowitz LE, Hariri S, Lin C, Dunne EF, Steinau M, McQuillan G, & Unger ER (2013) J Infect Dis 208, 385–393.

52. Hill MO (1973) Ecology 54, 427–432.

53. Jost L (2007) Ecology 88, 2427–2439.

